# A new SARS-CoV-2 lineage that shares mutations with known Variants of Concern is rejected by automated sequence repository quality control

**DOI:** 10.1101/2021.04.05.438352

**Authors:** Bryan Thornlow, Angie S. Hinrichs, Miten Jain, Namrita Dhillon, Scott La, Joshua D. Kapp, Ikenna Anigbogu, Molly Cassatt-Johnstone, Jakob McBroome, Maximilian Haeussler, Yatish Turakhia, Terren Chang, Hugh E Olsen, Jeremy Sanford, Michael Stone, Olena Vaske, Isabel Bjork, Mark Akeson, Beth Shapiro, David Haussler, A. Marm Kilpatrick, Russell Corbett-Detig

**Author notes:** co-first authors (Bryan Thornlow, Angie S. Hinrichs).

## Abstract

We report a SARS-CoV-2 lineage that shares N501Y, P681H, and other mutations with known variants of concern, such as B.1.1.7. This lineage, which we refer to as B.1.x (COG-UK sometimes references similar samples as B.1.324.1), is present in at least 20 states across the USA and in at least six countries. However, a large deletion causes the sequence to be automatically rejected from repositories, suggesting that the frequency of this new lineage is underestimated using public data. Recent dynamics based on 339 samples obtained in Santa Cruz County, CA, USA suggest that B.1.x may be increasing in frequency at a rate similar to that of B.1.1.7 in Southern California. At present the functional differences between this variant B.1.x and other circulating SARS-CoV-2 variants are unknown, and further studies on secondary attack rates, viral loads, immune evasion and/or disease severity are needed to determine if it poses a public health concern. Nonetheless, given what is known from well-studied circulating variants of concern, it seems unlikely that the lineage could pose larger concerns for human health than many already globally distributed lineages. Our work highlights a need for rapid turnaround time from sequence generation to submission and improved sequence quality control that removes submission bias. We identify promising paths toward this goal.

## Introduction

Rapid SARS-CoV-2 genome sequencing enables researchers to trace the virus’ evolution as it spreads and adapts within human populations. Several important mutations have arisen within the SARS-CoV-2 population that are thought to increase the transmissibility of the virus (Korber et al. 2020; Volz, Hill, et al. 2021; Volz, Mishra, et al. 2021) or to improve the virus’ ability to evade host immune defenses (Hoffmann et al. 2021; Planas et al. 2021). The early detection of new variant lineages of SARS-CoV-2 that might have similar traits is essential to prioritizing public health responses, developing variant-specific diagnostics and vaccines, and beginning research into the possible immunological and general health implications of newly discovered variants. Here, we report results from viral genome sequencing in Santa Cruz County, CA where we describe a newly discovered variant lineage, here denoted B.1.x in anticipation of a more refined classification later, that shares several mutations with known SARS-CoV-2 variants of concern. These mutations, combined with the recent spread of this variant, suggest that it warrants further study.

This is an ongoing study and we welcome any insights or collaborative efforts to improve our analysis. Please contact the authors if you would like to contribute to the study of B.1.x or to help our efforts to improve automated sequence quality control and rapid, unbiased reporting.

## Results and Discussion

### Sequence Data

We obtained a consensus sequence from 339 samples from Santa Cruz County. The majority of these samples are high quality, and 88% have a single maximally parsimonious placement on the global SARS-CoV-2 phylogeny that we maintain (Turakhia, Thornlow, et al. 2020). This suggests phylogenetic inference using these samples is reliable (Turakhia, Thornlow, et al. 2020). Approximately 58% of these 339 sequences are associated with lineages B.1.427 and B.1.429, first identified in California (Deng et al. 2021). We also detected two samples in our dataset of lineage B.1.1.7, first detected in the UK (Volz, Mishra, et al. 2021) and increasing in southern California, USA (Washington et al. 2021). We did not detect any other CDC-designated Variants of Concern (VOCs) or variants of interest (https://www.cdc.gov/coronavirus/2019-ncov/cases-updates/variant-surveillance/variant-info.html). The distribution of lineages in Santa Cruz County is similar to recent reports from the San Francisco Bay Area approximately 100km to the north (Peng et al. 2021) (Figure 1).

**Figure 1.**
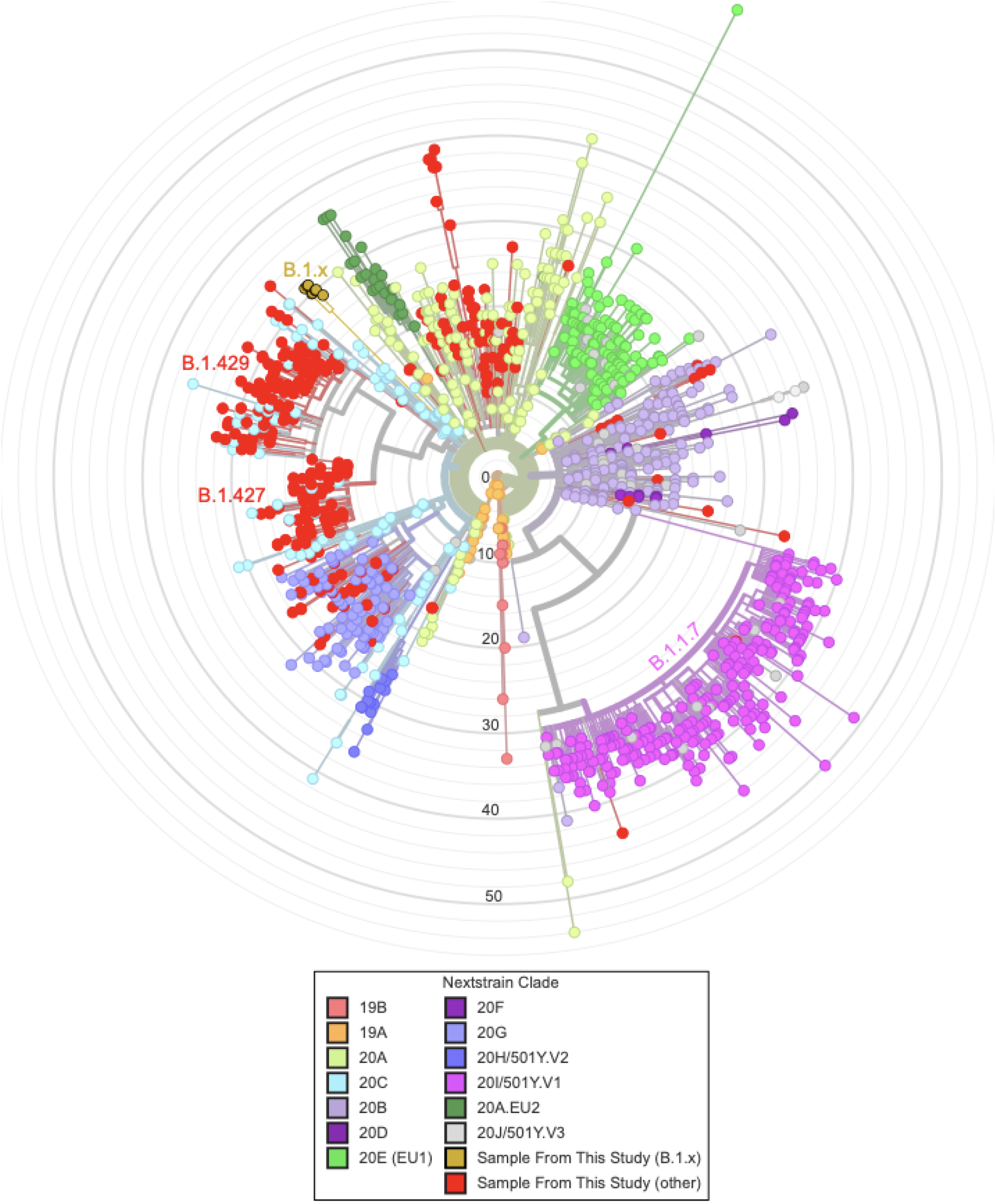
The phylogenetic distribution of 339 samples obtained from SARS-CoV-2 sequencing in Santa Cruz County plus 1000 samples from elsewhere. The tree is produced via the UShER web portal at hgPhyloPace (https://genome.ucsc.edu/cgi-bin/hgPhyloPlace). To produce it we added the 339 genomes from the Santa Cruz County samples to a global phylogeny of > 1 million SARS CoV-2 genomes and then pruned back to retain only the Santa Cruz genomes plus 1000 others selected at random. We visualized the tree using the Auspice.us platform. The 339 samples from Santa Cruz County are colored in red, with the eight samples representing B.1.x highlighted in gold, and the remaining 1000 samples are colored by Nextstrain clade. Note that clade sizes reflect both prevalence and local sampling effort and we have not attempted to correct for the effect of either.

### A new largely undescribed lineage

Eight samples collected between mid February and early March represent a largely undescribed lineage. Hereafter we refer to this as lineage B.1.x as it is nested within the prevalent lineage B.1 in our phylogeny (Nextstrain 20C). Using the pangolin tool (https://github.com/cov-lineages/pangolin/blob/master/github.com/cov-lineages/pangolin), our samples are assigned either B.1 or B.1.324 (Table S1, Rambaut et al. 2020). We note that some samples in this clade have been referred to as B.1.324.1 by COG-UK (https://assets.publishing.service.gov.uk/government/uploads/system/uploads/attachment_data/file/972247/Variants_of_Concern_VOC_Technical_Briefing_7_England.pdf) and the phylogenetic origins of this lineage are challenging to establish with certainty (see https://github.com/cov-lineages/pango-designation/issues/25, with analysis by A. Hinrichs).

### This lineage shares mutations with known VOCs

These samples contain several mutations shared with B.1.1.7 and other VOCs. The eight samples share six non-synonymous mutations within the S or Spike protein relative to the Wuhan-Hu-1 SARS-CoV-2 reference sequence (RefSeq NC_045512.2). Specifically, each sample contains Spike mutations S494P, N501Y, D614G, P681H, K854N, and E1111K. Among these, three mutations are suspected to have an effect on the viral fitness and transmissibility. Specifically, N501Y is thought to be important for viral replication because it enables the virus to bind ACE2 and enter host cells more efficiently (Gu et al. 2020; Starr et al. 2020). S494P is also located within the ACE2 receptor binding domain and experimental evidence suggests that mutations at this position decrease antibody binding affinity (Starr et al. 2020). Similarly, P681H is located within the spike protein furin cleavage site which is thought to be a hotspot of viral adaptive evolution (*e.g*., Peng et al. 2021; Zuckerman et al. 2021). D614G became globally dominant in 2020 possibly due to higher viral loads (Volz, Hill, et al. 2021). The effect of the other two Spike mutations is unknown.

In addition to the Spike mutations, B.1.x includes N:M234I (G28975A), which also appears in Variants of Interest B.1.526 and P.2 (G28975T). The three nucleotide mutations that cause N:M234I (G28975A, G28975C, G28975T) have all been observed at the roots of several Pango lineages (Table S2) and the frequency of N:M234I in 480,704 samples available from GISAID as of 2 April 2021 with collection dates 2021-01-01 to 2021-03-31 is 7.0%. N:M234I has been predicted to be stabilizing for the protein structure (Jacob et al. 2020). More generally, because each of these mutations appear to have occurred independently from other VOCs in B.1.x (see discussion of recombination below), these substitutions reveal significant evolutionary parallelism between B.1.x and known VOCs.

### Recent evolutionary dynamics of B.1.x

Samples from B.1.x comprise approximately 2.4% of the 339 samples that we sequenced from October through early April in Santa Cruz County, CA. We observed a striking increase rising from less than 1% in January to more than 10% by mid-March (Figure 2). The estimated rate of increase suggests that the number of B.1.x cases might be increasing over time despite total COVID-19 cases decreasing sharply during this period (Figure S1). However, the increase in frequency is uncertain because the variant was not detected in the 16 most recently collected samples, collected between March 19, 2021 and March 29, 2021 (Figure 2). The average rate of increase of this lineage was similar to the increase of B.1.1.7 in southern California, but with a slightly later rise in frequency (Figure S2). Further, the reconstructed phylogeny and case investigations of these samples indicate significant local transmission rather than several recurrent introductions into the Santa Cruz area. This suggests the variant B.1.x might be growing in frequency. This apparent increase in frequency within Santa Cruz County is noteworthy in light of the fact that B.1.1.7 has been introduced to the area at least twice, but has not increased in frequency as occurred in Southern California (Figure S2) and in other places worldwide *(e.g*. Washington et al. 2021).

**Figure 2.**
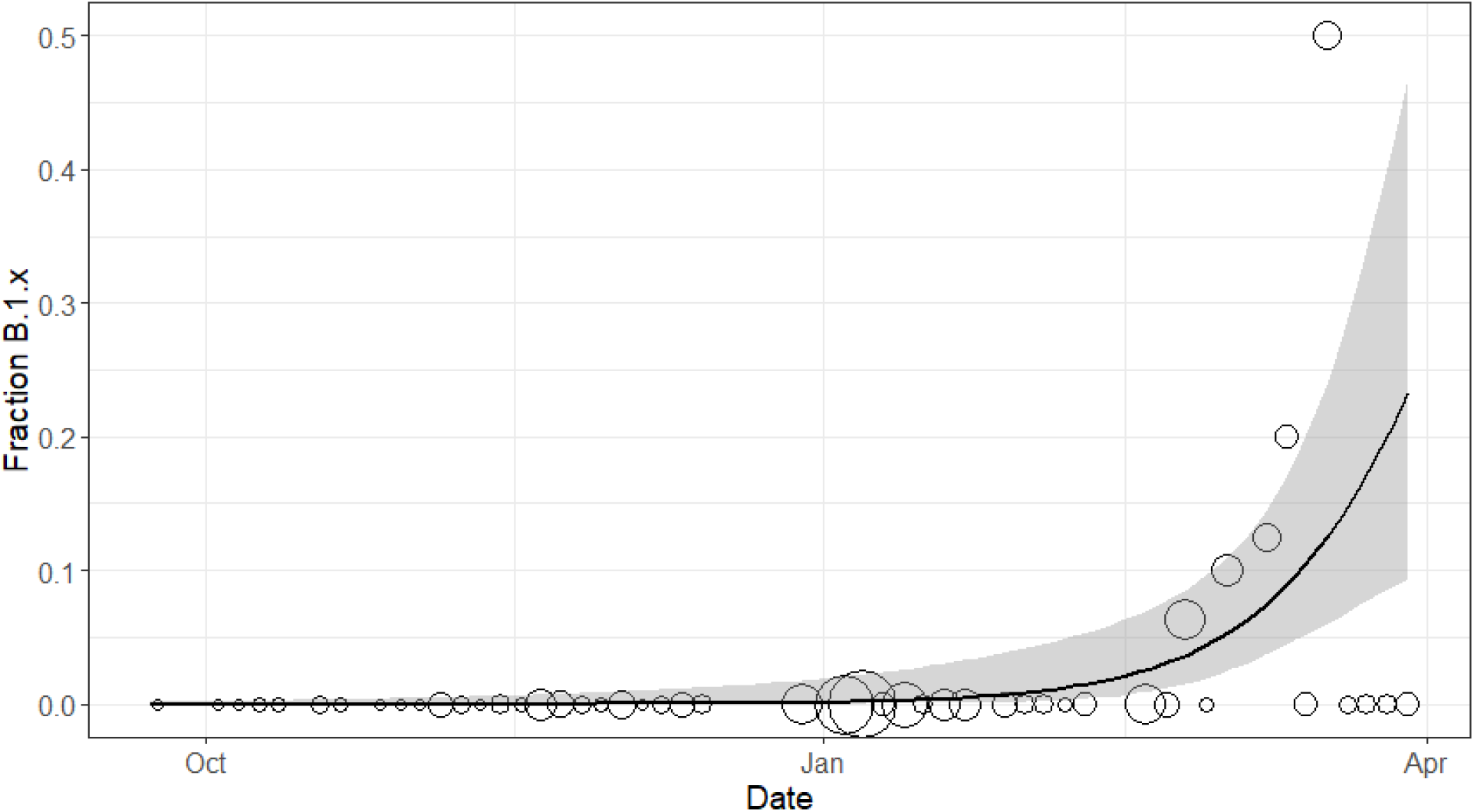
Increase in the fraction of samples from B.1.x over time. Points show the fraction of 339 samples from Santa Cruz County grouped into 3-day time intervals belonging to the B.1.x lineage, with the size of points showing the square root of the sample size (mean 6.7, range 1-48). The fitted line and shaded 95% confidence interval are based on a generalized linear model with a binomial distribution (intercept −1179 ±375; slope 0.063±0.020; P = 0.002).

We caution that there are important limitations to this study. (1) Our relatively small sample size imposes substantial uncertainty in our estimated frequencies, (2) we did not detect B.1.x in the most recent 16 samples from late March (which represent only a fraction of positive cases, see Table S3), and (3) our samples were taken opportunistically from positive tests and are not a random sample of infections or cases. The non-random sampling and underlying phylogeny imposes complex dependencies on the data that are not part of the simple logistic regression analysis (see similar caveats for early investigations of other variants of SARS-CoV-2, e.g. Volz, Hill, et al. 2021; Volz, Mishra, et al. 2021).

Comparison against publicly available samples from across the United States indicates the B.1.x lineage is present in several US states including New York, Florida, Georgia, and Indiana. Additionally, phylogenetic evidence suggests successful establishment and ongoing transmission in each of those states as well as others (Figure 4).

### B.1.x does not appear to be a recombinant

Despite the fact that this lineage shares several mutations with established VOCs, homoplastic substitution and not recombination is the most likely explanation. Recent work suggests that recombination has occurred in SARS-CoV-2 (VanInsberghe et al. 2021; Varabyou et al. 2020). Nonetheless, recombination should produce an extended stretch of sequence similarly between the donor and recipient lineage, and visual inspection of each VOC that shares at least one mutation with this lineage does not reveal any such tracts (Figure 3). Instead, we find the nucleotide substitution pattern is largely consistent with other lineages within Pango clade B.1 (Nextstrain 20C).

**Figure 3:**
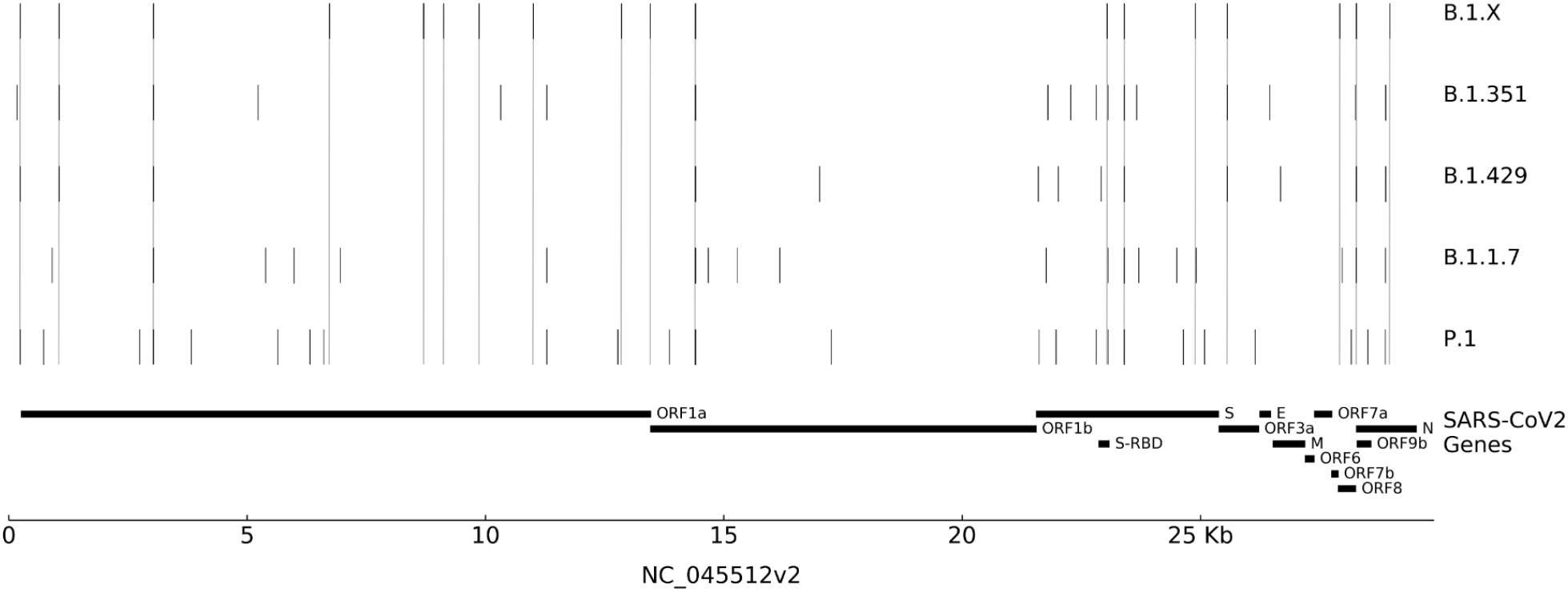
Sequence variation across each VOC and lineage B.1.x. Dashed lines indicate mutations relative to the reference sequence that are shared by B.1.x and at least one VOC. No stretch of variation appears to clearly match any single VOC, as we would expect if recent recombination generated the shared mutations. In particular, the spike region is quite distinct except for the shared mutations with known VOCs such as N501Y and P681H as noted in the main text.

**Figure 4:**
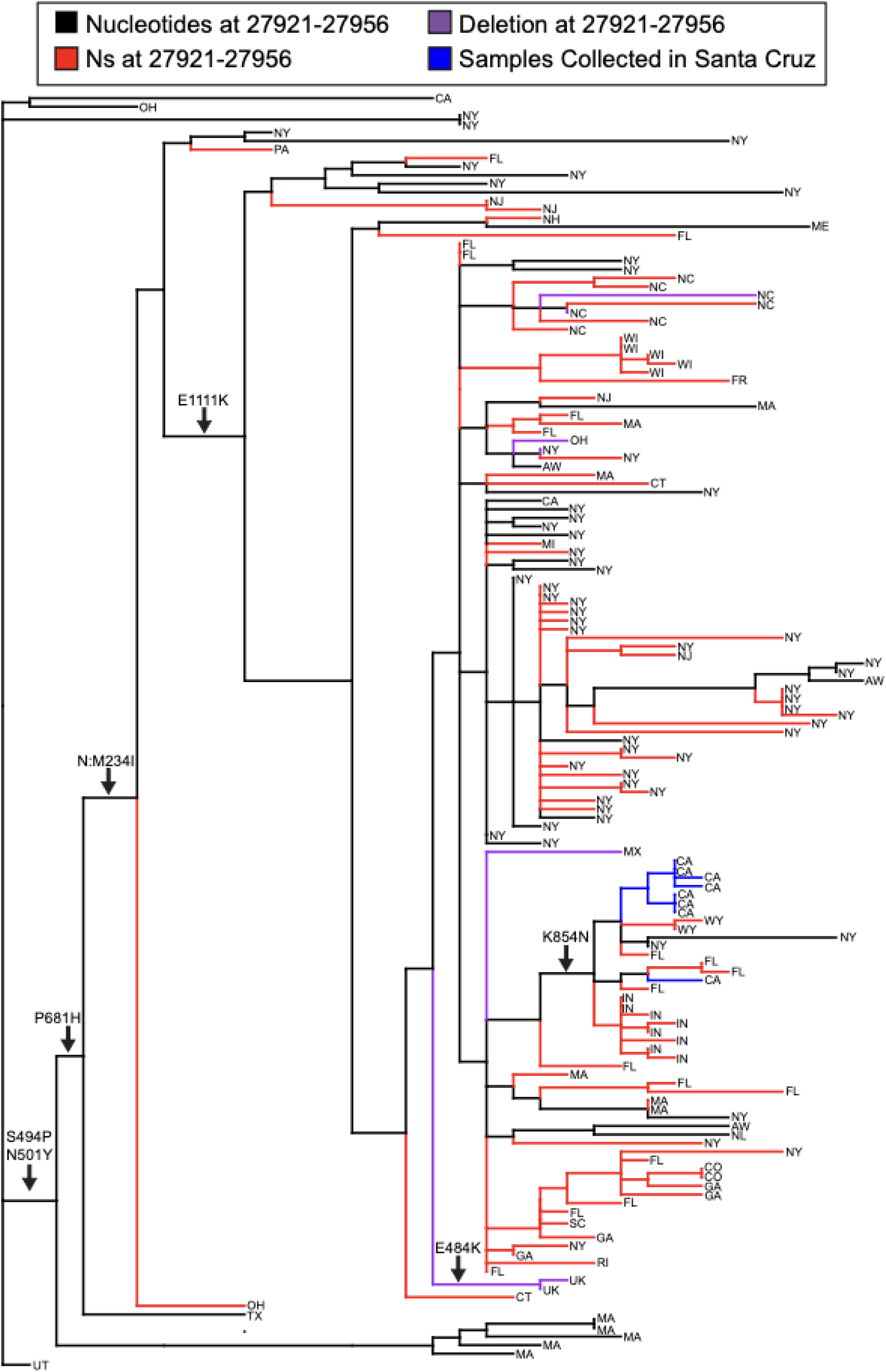
Phylogenetic distribution of 147 sequences, inferred based on whole-genome sequences and colored based on the region of a deletion of 35 bases in our samples of B.1.x and publicly available samples closely related to this clade. Our samples and others containing the deletion are shown in blue and purple respectively. Samples with missing data, N’s, throughout the deletion are shown in red, and samples with nucleotide sequence are shown in black. To accommodate slight variation in consensus sequences, we refer to all samples with at least 30 “-” characters in the multiple sequence alignment as deletion and similarly all sequences with at least 30 “N” characters as missing data. Branch lengths are proportional to the number of mutations inferred to occur on that branch. All spike amino-acid altering mutations in B.1.x are shown except D614G which occurred in an ancestor that is not shown in this phylogeny. Note that two samples isolated in the UK also contain E484K as indicated. The state or country abbreviation for each sample is shown at each leaf (UK = United Kingdom, MX = Mexico, FR = France, AW = Aruba, NL = Netherlands). The identifiers for the samples shown in this figure can be found in Table S4.

### B.1.x contains a large deletion in ORF8

B.1.x contains a 35bp deletion that induces a frameshift and early stop codon in ORF8 (bases 27922-27956 of RefSeq NC_045512.2) that is reminiscent of B.1.1.7, which also contains a nonsense mutation that causes a premature stop codon in ORF8. However, the functional significance of this mutation is not known. This parallel evolution with the B.1.1.7 lineage suggests that inactivation of ORF8 may be favorable to the virus, possibly in combination with shared amino acid substitutions within the Spike protein (Zinzula 2021).

Variable genome consensus generation methods and difficulties in accurately genotyping large indels make it challenging to establish the evolutionary pattern of this deletion with certainty. There appears to be dramatic variation in the presence/absence of this deletion across closely related sequences (Figure 4). However, many related consensus sequences have this section of the genome replaced with N’s, and others contain nucleotide sequences that suggest that the deletion may have been replaced with reference alleles (see below). We believe that it is unlikely that this large, specific deletion evolved more than once in this lineage. Conversely, a reversion to the reference, via reinsertion of identical 35bp nucleotide sequence is unlikely. Instead, we suggest that this highly unusual evolutionary pattern primarily reflects apparent differences in consensus genome assembly and submission methods where some tools may insert reference bases and others call deletions as missing data. Of course, confident phylogenetic inference with SARS-CoV-2 is difficult (Morel et al. 2020; Turakhia, De Maio, et al. 2020), so it is possible that errors in the inferred phylogeny contribute to this apparent improbable evolutionary pattern at this deletion. Comparison to raw sequence data will be necessary to confirm or refute this. However the pattern suggests that the deletion is a shared feature among most genomes in this lineage and that its apparent sporadic absence in some genomes is an artefact of the way virus genomes are currently assembled and submitted.

### Automated sequence QC tools reject B.1.x submission to databases

In submitting our sequences, we found that all eight genomes corresponding to the B.1.x lineage were initially rejected by both GISAID and Genbank due to the frameshift-inducing deletion of 35 bases in ORF8. To get them accepted required a few iterations of manual curation and sequence confirmation. An informal poll of other groups submitting sequences to these databases recently indicated that many are too busy to engage in this process and are setting aside genomes that do not initially pass automated sequence checks. This means it is possible that sequences belonging to this lineage are underreported in GISAID and Genbank. Although such automated checks are clearly essential for maintaining high quality sequence databases, they may also result in a significant bias in the SARS-CoV-2 variants that are present in the database. We suggest that rapid phylogenetic placement (using *e.g*., UShER, (Turakhia, Thornlow, et al. 2020)) during sequence submission might allow closely related sequences with novel variation to corroborate each other during batch submissions and across independent sequences, thus alleviating some of this submission burden. It is critical that these issues with GISAID and Genbank submission systems be improved, as they could hamper our ability to recognize and evaluate new variants of concern as they arise.

## Conclusion

Pathogen genomic surveillance has the potential to identify novel lineages with important immunological and epidemiological consequences. Our sequencing efforts have uncovered a potentially important new lineage, referred to as B.1.x here. Future work is clearly needed to determine whether B.1.x is increasing in frequency or spreading spatially in the United States or elsewhere. We emphasize that at present, while preliminary data shows evidence of an increase in frequency, the numbers are small, and direct evidence for enhanced transmissibility or immune evasion of this lineage, or differences in disease severity, is absent. Nonetheless, given the genomic parallels to known VOCs, further study of secondary attack rates, viral loads, antibody neutralization and disease severity of B.1.x and co-circulating variants would be prudent. If this variant is indeed underreported, then its prevalence may already be higher than is apparent.

Given what is known about currently circulating variants of concern, we do not expect B.1.x to become globally dominant. In particular, none of the isolates except for two from the UK contain E484K, which is present in other VOCs P.1, B.1.351 and in sublineages of B.1.1.7. This mutation is believed to be a key component of VOCs’ ability to evade the immune response (Collier et al. 2021; Resende et al. 2021; Nonaka et al. 2021). However, two B.1.x sequences isolated from travelers returning to the UK also contain E484K, and the effect of S494P is not as well characterized. If this sublineage expands, or if other sublineages secondarily acquire E484K or other mutations that enhance viral fitness, B.1.x may become more likely to impose a global health threat.

Our results highlight the need for expanded scrutiny of viral genomic data and improved quality control processes. Because sequences of B.1.x that accurately represent the deletion in ORF8 are rejected by automated quality checks, many may not be present in public databases. This difficulty blunts our efforts to carefully track the abundance and spread of this lineage across the USA and worldwide, and may do so for other new lineages. This work is not intended as a criticism of database quality control at GISAID or NCBI. Both are commendable, essential and significant efforts. However, improved tools that use similarity to accepted curated sequences and other features to more rapidly accept newly discovered variants will ease the burden on genome-submitting laboratories, and may be essential to timely and accurate tracing of the evolution of pathogens.

## Methods

### Sample Collection, Sequencing, and Assembly

Amplicon sequencing was performed using residual anonymized SARS-CoV-2 positive clinical samples. Viral RNA was extracted from residual anterior nares swabs collected in DNA/RNA Shield (ZymoResearch) using anion exchange chromatography on magnetic beads (ZymoResearch). Eluted RNA was re-analyzed by RT-qPCR using the CII Triplex SARS-CoV-2 Assay (EUA200510). RNA samples with Cycle Thresholds (Cts) for SARS-CoV-2 targets less than 30 were selected for amplicon sequencing.

We used Illumina’s COVIDSeq Test (RUO) (https://www.illumina.com/products/by-type/clinical-research-products/covidseq.html) workflow for generating SARS-CoV-2 sequencing data. Briefly this process involved the following steps: 1) The UCSC Molecular Diagnostics Lab re-isolated RNA for SARS-CoV-2 positive samples; 2) the RNA samples underwent reverse transcription and PCR amplification using manufacturer recommendations; 3) the amplicons were prepared into sequencing libraries using a bead-linked transposome tagmentation method, following the manufacturer’s recommendations; 4) the libraries were sequenced on the Illumina NextSeq550 using Mid Output v2.5 (150 cycle) sequencing kits, which generates 2×73 paired reads; 5) we generated consensus sequences for each sample using the Illumina DRAGEN COVIDSeq Test RUO v1.3.0 application available in BaseSpace using the default parameters.

### Assembly Confirmation

We manually confirmed the quality of our genome assemblies in the region of the large deletion. Short read alignments appear consistent with the 35bp deletion as reported in our sequences and are confirmed by several others in the GISAID database that are tagged as having received manual curation efforts.

### Primary Analysis

We identified closely related samples and genetic variation present in each newly sequenced genome by uploading the consensus sequences to the UShER web interface (https://genome.ucsc.edu/cgi-bin/hgPhyloPlace, (Turakhia, Thornlow, et al. 2020)). This interface automatically generates the subtree of the most closely related genomes in a format compatible with the Auspice visualization platform (Hadfield et al. 2018). Additionally, we use the “view in Genome Browser” option to manually scan sequences for mutations from known VOCs and for other mutations in similar genomic positions within the Spike protein within the SARS-CoV-2 Genome Browser (Fernandes et al. 2020). The system was also used to extract 1000 random genomes from the tree to visualize newly sequenced genomes on the background of SARS-CoV-2 genomic variation.

### Phylogenetic Inference

We used the UShER web portal as our primary phylogenetic inference framework (Turakhia, Thornlow, et al. 2020). This is described previously and is a powerful way to identify transmission among the most closely related samples.

To confirm that the tree used in these analyses is the optimal or nearly optimal phylogeny given our sequencing data, we optimized the tree using IQ-TREE 2 (Minh et al. 2020). We used the following command: *iqtree2 -s mainClade.pruned.fa -m JC+I+R5 -fast -nt 4 -optalg 1-BFGS -experimental --suppress-list-of-sequences -t NJ-R --no-opt-gamma-inv -blmin 0.0000000001 -pre mainClade.pruned.iqtree2* where the input fasta was pruned to only the 147 samples shown in Figure 4. We then compared the tree output by this command to our base tree, finding few differences. The two trees have a Robinson-Foulds distance of 66.0 (Bogdanowicz, Giaro, and Wróbel 2012; Robinson and Foulds 1981).

### Statistical analyses

We analyzed temporal variation in the frequency of the B.1.x variant with a generalized linear model with a binomial distribution and a logit link. We used the date that each sample was collected as the predictor. We analyzed temporal variation in the frequency of the B.1.1.7 variant using data from the Helix® COVID-19 Surveillance Dashboard (Helix.com/covid19db and https://raw.githubusercontent.com/myhelix/helix-covid19db/master/counts_by_state.csv accessed on April 3, 2021) which includes samples primarily from southern California and mostly from San Diego County (Nicole Washington, Helix, personal communication). We analyzed the fraction of all samples that exhibited S-gene target failure (SGTF) from December 3, 2020 to March 29, 2021. During this period 445/532 or 84% of SGTF samples that were sequenced were B.1.1.7.

## Supporting information

Figures S1 and S2, Tables S2 and S3, all supplemental figure and table legends

## Acknowledgements

The authors thank David Ghilarducci, Cal Gordon, Emily Chung, Gail Newel, Ramy Husseini and Mikala Caton with the Santa Cruz Department of Public Health, Deborah Wadford and Joel Sevinsky with the California Department of Public Health, as well as Duncan MacCannell, Ryan Tewhey and the entire CDC-SPHERES community. We are also indebted to the outstanding efforts of the GISAID and Genbank database teams and all the laboratories that submit data to them (Table S5).

## Data Availability

All sequence data produced in this work is available from GISAID and Genbank with accession numbers indicated in Table S1.

