## Supplementary material for "A new SARS-CoV-2 lineage that shares mutations with known Variants of Concern is rejected by automated sequence repository quality control": Table S5

We gratefully acknowledge the following Authors from the Originating laboratories responsible for obtaining the specimens, as well as the Submitting laboratories where the genome data were generated and shared via GISAID, on which this research is based.

All Submitters of data may be contacted directly via www.gisaid.org

Authors are sorted alphabetically.

| Accession ID | Originating Laboratory | Submitting Laboratory | Authors |
|---|---|---|---|
| EPI_ISL_1014240 | Dutch COVID-19 response team | National Institute for Public Health and the Environment (RIVM) | Adam Meijer, Harry Vennema, Dirk Eggink, Jeroen Cremer, Sharon van den Brink, Bas van der Veer, AnneMarie van den Brandt, Florian Zwagemaker, Dennis Schmitz, Chantal Reusken, on behalf of the national COVID-19 response team |
| EPI_ISL_1016170 | NYU Langone Health | Departments of Pathology and Medicine, New York University School of Medicine | Adriana Heguy, Dacia Dimartino, Emily Guzman, Christian Marier, Peter Meyn, Sitharam Ramaswami, Gael Westby, Paul Zappile, Yutong Zhang, Paolo Cotzia, Guiqing Wang |
| EPI_ISL_1016573 | Helix/Illumina | Respiratory Viruses Branch, Division of Viral Diseases, Centers for Disease Control and Prevention | Peter W. Cook, Dakota Howard, Dhwani Batra, Ben L. Rambo-Martin, Eileen de Feo, Jan Antico, Christine Tran, Matthew Tolentino, Shannon Wickline, Kim Gietzen, Brad Sickler, Jingtao Liu, Eric Allen, Phil Febbo, Summer Galloway, Nicole L. Washington, Simon White, Geraint Levan, Kelly Schiabor Barrett, Elizabeth Cirulli, Alexandre Bolze, Ary Ascencio, Charlotte Rivera-Garcia, Ryan Cho, Jason Nguyen, Sherry Wang, Jimmy Ramirez, Tyler Cassens, Efren Sandoval, Magnus Isaksson, William Lee, David Becker, Marc Laurent, James Lu, Clinton R. Paden, Suxiang Tong, Duncan MacCannell |
| EPI_ISL_1017816, EPI_ISL_1017822 | University of Wisconsin-Madison AIDS Vaccine Research Laboratories | University of Wisconsin-Madison AIDS Vaccine Research Laboratories | Gage Moreno, Katarina Braun, et al. AIDS Vaccine Research Laboratories |
| EPI_ISL_1021530, EPI_ISL_1021865, EPI_ISL_1021888 | Laboratory Corporation of America | Respiratory Viruses Branch, Division of Viral Diseases, Centers for Disease Control and Prevention | Peter W. Cook, Dakota Howard, Dhwani Batra, Ben L. Rambo-Martin, Clinton R. Paden, Suxiang Tong, Duncan MacCannell |
| EPI_ISL_1027033 | Fulgent Genetics | Fulgent Genetics | Harry Gao, Mickey Li, John Gao, Joseph Fierro, Benafsh Sapra, Becky Tsai, Yan Meng, Doreen Ng, James Xie |
| EPI_ISL_1031320, EPI_ISL_1037713 | Laboratory Corporation of America | Respiratory Viruses Branch, Division of Viral Diseases, Centers for Disease Control and Prevention | Peter W. Cook, Dakota Howard, Dhwani Batra, Ben L. Rambo-Martin, Clinton R. Paden, Suxiang Tong, Duncan MacCannell |
| EPI_ISL_1039916 | University of Wisconsin-Madison AIDS Vaccine Research Laboratories | University of Wisconsin-Madison AIDS Vaccine Research Laboratories | Gage Moreno, Katarina Braun, et al. AIDS Vaccine Research Laboratories |
| EPI_ISL_1041259, EPI_ISL_1041366, EPI_ISL_1041389, EPI_ISL_1041646 | Pandemic Response Lab - NYC | Pandemic Response Lab, R&D | Henry Lee, Michael Hammerling, Melissa Hopkins, Cybill del Castillo, William Ward, Pradeep Bugga, Haiping Hao, Jon Laurent |
| EPI_ISL_1087279 | NYU Langone Health | Departments of Pathology and Medicine, New York University School of Medicine | Adriana Heguy, Dacia Dimartino, Emily Guzman, Christian Marier, Peter Meyn, Sitharam Ramaswami, Gael Westby, Paul Zappile, Yutong Zhang, Paolo Cotzia, Guiqing Wang |
| EPI_ISL_1089592 | Dutch COVID-19 response team | National Institute for Public Health and the Environment (RIVM) | Adam Meijer, Harry Vennema, Dirk Eggink, Jeroen Cremer, Sharon van den Brink, Bas van der Veer, AnneMarie van den Brandt, Florian Zwagemaker, Dennis Schmitz, Chantal Reusken, on behalf of the national COVID-19 response team |
| EPI_ISL_1097808, EPI_ISL_1097811, EPI_ISL_1097904, EPI_ISL_1097961, EPI_ISL_1097971, EPI_ISL_1097972, EPI_ISL_1097983, EPI_ISL_1098189, EPI_ISL_1098318, EPI_ISL_1098361, EPI_ISL_1098549, EPI_ISL_1098585 | | | |
| see above | Pandemic Response Lab - NYC | Pandemic Response Lab, R&D | Henry Lee, Michael Hammerling, Melissa Hopkins, Cybill del Castillo, Shinyoung Clair Kang, William Ward, Pradeep Bugga, Haiping Hao, Jon Laurent |
| EPI_ISL_1132137 | Florida Bureau of Public Health Laboratories | Florida Bureau of Public Health Laboratories | Sarah Schmedes, Jason Blanton |
| EPI_ISL_1134507 | Helix/Illumina | Respiratory Viruses Branch, Division of Viral Diseases, Centers for Disease Control and Prevention | Peter W. Cook, Dakota Howard, Dhwani Batra, Ben L. Rambo-Martin, Eileen de Feo, Jan Antico, Christine Tran, Matthew Tolentino, Shannon Wickline, Kim Gietzen, Brad Sickler, Jingtao Liu, Eric Allen, Phil Febbo, Summer Galloway, Nicole L. Washington, Simon White, Geraint Levan, Kelly Schiabor Barrett, Elizabeth Cirulli, Alexandre Bolze, Ary Ascencio, Charlotte Rivera-Garcia, Ryan Cho, Jason Nguyen, Sherry Wang, Jimmy Ramirez, Tyler Cassens, Efren Sandoval, Magnus Isaksson, William Lee, David Becker, Marc Laurent, James Lu, Clinton R. Paden, Suxiang Tong, Duncan MacCannell |
| EPI_ISL_1160540, EPI_ISL_1160617, EPI_ISL_1160804, EPI_ISL_1162256, EPI_ISL_1162338, EPI_ISL_1162341, EPI_ISL_1162371, EPI_ISL_1162450, EPI_ISL_1162595 | Laboratory Corporation of America | Respiratory Viruses Branch, Division of Viral Diseases, Centers for Disease Control and Prevention | Peter W. Cook, Dakota Howard, Dhwani Batra, Ben L. Rambo-Martin, Minoo Agarwal, Eyad Almasri Debbie Boles, Ayla Burns, Nuthawin Charoensri, Oren Cohen, Susan Countryman, Mary Ann Cristobal, Bobbi Croy, Suzanne Dale, Hrushikesh Deshmukh, Amanda Douglas, Vincent Drouillon, Marcia Eisenberg, Howard Engler, Rama Ghatti, Prashant Gupta, Susan Hicks, Jake Humphrey, Lax Iyer, Manoj Jain, Mohan Kolli, Brian Krueger, Tim Kuphal, Stanley Letovsky, Michael Levandoski, Craig Lukasik, Jonathan Meltzer, Brian Norvell, Mindy Nye, Scott Parker, Christos Petropoulos, John Pruitt, Steven Ragan, Scott Ryan, Mike Sapeta, Jana Schroth, Suresh Babu Selvaraju, Goran Stevovic, Amanda Suchanek, Andrea Throop, Lyndon Tilson, Thomas Urban, Joe Voshell, Kimberly Wagner, Jonathan Williams, Mary Williamson, Qian Zeng, Tricia Zwiefelhofer, Clinton R. Paden, Suxiang Tong, Duncan MacCannell |
| EPI_ISL_1163630 | Maine HETL | Tewhey Lab, The Jackson Laboratory | Matluk,N., Dewey,H., Iosue,F., Barter,M., Lynch,R., Munger,H. and Tewhey,R. |
| EPI_ISL_1171856 | MSHS Clinical Microbiology Laboratories | MSHS Pathogen Surveillance Program | Fatima Amanat, Mahima Thapa, Tinting Lei, Shaza M. Sayed Ahmed, Daniel C. Adelsberg, Juan Manuel Carreno, Shirin Strohmeier, Aaron J. Schmitz, Sarah Zafar, Julian Q Zhou, Willemijn Rijnink, Hala Alshammary, Nicholas Borcherding, Ana Gonzalez Reiche, Komal Srivastava, Emilia Mia Sordillo, Harm van Bakel, The Personalized Virology Initiative, Jackson S. Turner, Goran Bajic, Viviana Simon, Ali H. Ellebedy, Florian Krammer |
| EPI_ISL_1171857 | Department of Microbiology, Icahn School of Medicine at Mount Sinai | MSHS Pathogen Surveillance Program | Fatima Amanat, Mahima Thapa, Tinting Lei, Shaza M. Sayed Ahmed, Daniel C. Adelsberg, Juan Manuel Carreno, Shirin Strohmeier, Aaron J. Schmitz, Sarah Zafar, Julian Q Zhou, Willemijn Rijnink, Hala Alshammary, Nicholas Borcherding, Ana Gonzalez Reiche, Komal Srivastava, Emilia Mia Sordillo, Harm van Bakel, The Personalized Virology Initiative, Jackson S. Turner, Goran Bajic, Viviana Simon, Ali H. Ellebedy, Florian Krammer |
| EPI_ISL_1172465, EPI_ISL_1172549, EPI_ISL_1172579, EPI_ISL_1172666, EPI_ISL_1172674, EPI_ISL_1172707, EPI_ISL_1172788, EPI_ISL_1172891, EPI_ISL_1172967, EPI_ISL_1173069 | Pandemic Response Lab - NYC | Pandemic Response Lab, R&D | Henry Lee, Michael Hammerling, Melissa Hopkins, Cybill del Castillo, Shinyoung Clair Kang, William Ward, Pradeep Bugga, Haiping Hao, Jon Laurent |
| EPI_ISL_1182017 | Yale Clinical Virology Lab | Grubaugh Lab - Yale School of Public Health | Joseph Fauver, Mallery Breban, Isabell Ott, Tara Alpert, Mary Petrone, Anderson Brito, Chantal Vogels, Annie Watkins, Chaney Kalinich, Marie L. Landry, Nathan Grubaugh |
| EPI_ISL_1184172, EPI_ISL_1184195, EPI_ISL_1184202 | Indiana Animal Disease Diagnostic Laboratory | Carpi Laboratory - Purdue University | Jack Dorman, Ilinca I Ciubotariu, Lev Gorenstein, Abebe A Fola, G Kenitra Hendrix, Rebecca P Wilkes, Giovanna Carpi |
| EPI_ISL_1191477, EPI_ISL_1191535, EPI_ISL_1191536 | Florida Bureau of Public Health Laboratories | Florida Bureau of Public Health Laboratories | Sarah Schmedes, Jason Blanton |
| EPI_ISL_1195883 | NYU Langone Health | Departments of Pathology and Medicine, New York University School of Medicine | Adriana Heguy, Dacia Dimartino, Emily Guzman, Christian Marier, Peter Meyn, Sitharam Ramaswami, Gael Westby, Paul Zappile, Yutong Zhang, Paolo Cotzia, Guiqing Wang |
| EPI_ISL_1202566 | Ohio State University Wexner Medical Center | James Molecular Laboratory | Jones D, Chang Y-S, Ru P, Chappell D, Snyder P, Koenig S, Corcoran S, Pancholi P |
| EPI_ISL_1218767, EPI_ISL_1218778, EPI_ISL_1218794, EPI_ISL_1218810, EPI_ISL_1218932 | Florida Bureau of Public Health Laboratories | Florida Bureau of Public Health Laboratories | Sarah Schmedes, Jason Blanton |
| EPI_ISL_1233371 | Massachusetts State Public Health Laboratory | Massachusetts State Public Health Laboratory | Andrew Lang, Timelia Fink, Glen Gallagher, Sandra Smole |

| | | | |
|---|---|---|---|
| EPI_ISL_1239848 | New York Presbetyrian / Weill Cornell Medicine | Grubaugh Lab - Yale School of Public Health | Mary Petrone, Joseph Fauver, Tara Alpert, Anderson Brito, Mallery Breban, Anne Wyllie, Chantal Vogels, Annie Watkins, Chaney Kalinich, Isabel Ott, Nathan Grubaugh |
| EPI_ISL_1239898, EPI_ISL_1239919 | Florida Bureau of Public Health Laboratories | Florida Bureau of Public Health Laboratories | Sarah Schmedes, Jason Blanton |
| EPI_ISL_1240758 | University Hospitals Translational Laboratory (UHTL), University Hospitals | University Hospitals Translational Laboratory (UHTL), University Hospitals | Sadri,N., Alouani,D., Song,X. |
| EPI_ISL_1240807, EPI_ISL_1240862 | Clinical Molecular Microbiology Laboratory, UNC Hospitals | Jeremy Wang | Jeremy Wang, Alexander Rubinsteyn, Colleen Rice, Jason Smedberg, Shawn Hawken, Melissa Miller, Corbin Jones, Robert Hagan |
| EPI_ISL_1241883 | CENTRE HOSPITALIER EDMOND GARCIN | CNR Virus des Infections Respiratoires - France SUD | Antonin Bal, Gregory Destras, Gwendolyne Burfin, Hadrien Regue, Quentin Semanas, Martine Valette, Bruno Lina, Laurence Josset |
| EPI_ISL_1250418 | Michigan Department of Health and Human Services, Bureau of Laboratories | Michigan Department of Health and Human Services, Bureau of Laboratories | Blankenship HM, Riner D, Soehnlen MK |
| EPI_ISL_1252347 | University of Wisconsin-Madison AIDS Vaccine Research Laboratories | University of Wisconsin-Madison AIDS Vaccine Research Laboratories | Gage Moreno, Katarina Braun, et al. AIDS Vaccine Research Laboratories |
| EPI_ISL_1255279, EPI_ISL_1255280, EPI_ISL_1255282, EPI_ISL_1255285, EPI_ISL_1255289 | Indiana Animal Disease Diagnostic Laboratory | Carpi Laboratory - Purdue University | Jack Dorman, Ilinca I Ciubotariu, Nicole M Perry, Jobin J Kattoor, Lev Gorenstein, Abebe A Fola, G Kenitra Hendrix, Rebecca P Wilkes, Giovanna Carpi |
| EPI_ISL_1258557, EPI_ISL_1258558, EPI_ISL_1258559, EPI_ISL_1258560, EPI_ISL_1258977 | Pandemic Response Lab - NYC | Pandemic Response Lab, R&D | Henry Lee, Michael Hammerling, Melissa Hopkins, Cybill del Castillo, Shinyoung Clair Kang, William Ward, Pradeep Bugga, Haiping Hao, Jon Laurent |
| EPI_ISL_1288297 | Laboratorio Central de Epidemiología (LCE) | Instituto de Biotecnología de la UNAM | Consorcio Mexicano de Vigilancia Genómica (CoViGen-Mex). Authors (in alphabetical order): Julio Elias Alvarado-Yaah, Carlos F. Arias, Santiago Ávila-Ríos, Víctor Hugo Borja-Aburto, Celia Boukadida, Juan Bautista Chale-Dzul , José Antonio Enciso-Moreno, Gloria Elena Espinoza-Ayala, Fernando Fontove-Herrera, Concepción Grajales-Muñiz, Ricardo Grande, Alfredo Herrera-Estrella, Carla Ivón Herrera-Najera, Pavel Isa, Brenda Irasema Maldonado-Meza, Bernardo Martínez-Miguel, Margarita Matías-Florentino, María Guadalupe de Jesús Mireles-Rivera, Gloria María Molina-Salinas, Hector Montoya-Fuentes, José Esteban Muñoz-Medina, José de Jesús Nuñez-Contreras, Alicia Ocaña-Mondragón, Luis Alberto Ochoa-Carrera, Hector Esteban Paz-Juárez, Francisco Pulido, Helen Haydee Fernanda Ramírez-Plascencia, Angel Gustavo Salas-Lais, Jorge Ivan Salinal-Nevarez, Alejandro Sanchez-Flores, Clara Esperanza Santacruz-Tinoco, María Guadalupe Santiago-Mauricio , Nelly Sélem-Mojica, Blanca Taboada , Gloria Vazquez |
| EPI_ISL_1289136 | Dutch COVID-19 response team | National Institute for Public Health and the Environment (RIVM) | Adam Meijer, Harry Vennema, Dirk Eggink, Jeroen Cremer, Sharon van den Brink, Bas van der Veer, AnneMarie van den Brandt, Florian Zwagemaker, Dennis Schmitz, Chantal Reusken, on behalf of the national COVID-19 response team |
| EPI_ISL_1303675 | Houston Methodist Hospital | Houston Methodist Hospital | S. Wesley Long, Randall J. Olsen, Paul A. Christensen, Sishir Subedi, Robert Olson, James J. Davis, Matthew Ojeda Saavedra, Prasanti Yerramilli, Layne Pruitt, Kristina Reppond, Madison N. Shyer, Jessica Cambric, Ilya J. Finkelstein, Jimmy Gollihar, and James M. Musser |
| EPI_ISL_1306258, EPI_ISL_1306329, EPI_ISL_1306441, EPI_ISL_1306716, EPI_ISL_1306773, EPI_ISL_1306804, EPI_ISL_1307232, EPI_ISL_1307507, EPI_ISL_1307508, EPI_ISL_1307618 | Pandemic Response Lab - NYC | Pandemic Response Lab, R&D | Henry Lee, Michael Hammerling, Melissa Hopkins, Cybill del Castillo, Shinyoung Clair Kang, William Ward, Pradeep Bugga, Sol Rey, Dylan Law, Haiping Hao, Jon Laurent |
| EPI_ISL_1314235 | University of Wisconsin-Madison AIDS Vaccine Research Laboratories | University of Wisconsin-Madison AIDS Vaccine Research Laboratories | Gage Moreno, Katarina Braun, et al. AIDS Vaccine Research Laboratories |
| EPI_ISL_1319338, EPI_ISL_1319575, EPI_ISL_1319630, EPI_ISL_1319733, EPI_ISL_1320413, EPI_ISL_1320597, EPI_ISL_1320664, EPI_ISL_1320913, EPI_ISL_1320955, EPI_ISL_1320958 | Laboratory Corporation of America | Centers for Disease Control and Prevention Division of Viral Diseases, Pathogen Discovery | Peter W. Cook, Dakota Howard, Dhwani Batra, Ben L. Rambo-Martin, Minoo Agarwal, Eyad Almasri, Debbie Boles, Ayla Burns, Nuthawin Charoensri, Oren Cohen, Susan Countryman, Mary Ann Cristobal, Bobbi Croy, Suzanne Dale, Hrushikesh Deshmukh, Amanda Douglas, Vincent Drouillon, Marcia Eisenberg, Howard Engler, Rama Ghatti, Prashant Gupta, Susan Hicks, Jake Humphrey, Lax Iyer, Manoj Jain, Mohan Kolli, Brian Krueger, Tim Kuphal, Stanley Letovsky, Michael Levandoski, Craig Lukasik, Jonathan Meltzer, Brian Norvell, Mindy Nye, Scott Parker, Christos Petropoulos, John Pruitt, Steven Ragan, Scott Ryan, Mike Sapeta, Jana Schroth, Suresh Babu Selvaraju, Goran Stevovic, Amanda Suchanek, Andrea Throop, Lyndon Tilson, Thomas Urban, Joe Voshell, Kimberly Wagner, Jonathan Williams, Mary Williamson, Qian Zeng, Tricia Zwiefelhofer, Clinton R. Paden, Suxiang Tong, Duncan MacCannell |
| EPI_ISL_1321682 | Clinical Molecular Microbiology Laboratory, UNC Hospitals | Jeremy Wang | Jeremy Wang, Alexander Rubinsteyn, Colleen Rice, Jason Smedberg, Shawn Hawken, Melissa Miller, Corbin Jones, Robert Hagan |
| EPI_ISL_1322796 | NYU Langone Health | Departments of Pathology and Medicine, New York University School of Medicine | Adriana Heguy, Dacia Dimartino, Emily Guzman, Christian Marier, Peter Meyn, Sitharam Ramaswami, Gael Westby, Paul Zappile, Yutong Zhang, Paolo Cotzia, Guiqing Wang |
| EPI_ISL_1337613, EPI_ISL_1337614 | Wyoming Public Health Laboratory | Wyoming Public Health Laboratory | Jim Mildenberger, Wanda Manley, Noah Hull, Taylor Fearing, Lynette Gumbleton, Channing Weber, Ashley Norberg, Chayse Rowley, Marley Goetz, Brian Dominguez, Elliot Thomasson, Cari Sloma, and Rob Christensen |
| EPI_ISL_1338462, EPI_ISL_1338621, EPI_ISL_1338699, EPI_ISL_1338885 | Laboratory Corporation of America | Centers for Disease Control and Prevention Division of Viral Diseases, Pathogen Discovery | Peter W. Cook, Dakota Howard, Dhwani Batra, Ben L. Rambo-Martin, Minoo Agarwal, Eyad Almasri, Debbie Boles, Ayla Burns, Nuthawin Charoensri, Oren Cohen, Susan Countryman, Mary Ann Cristobal, Bobbi Croy, Suzanne Dale, Hrushikesh Deshmukh, Amanda Douglas, Vincent Drouillon, Marcia Eisenberg, Howard Engler, Rama Ghatti, Prashant Gupta, Susan Hicks, Jake Humphrey, Lax Iyer, Manoj Jain, Mohan Kolli, Brian Krueger, Tim Kuphal, Stanley Letovsky, Michael Levandoski, Craig Lukasik, Jonathan Meltzer, Brian Norvell, Mindy Nye, Scott Parker, Christos Petropoulos, John Pruitt, Steven Ragan, Scott Ryan, Mike Sapeta, Jana Schroth, Suresh Babu Selvaraju, Goran Stevovic, Amanda Suchanek, Andrea Throop, Lyndon Tilson, Thomas Urban, Joe Voshell, Kimberly Wagner, Jonathan Williams, Mary Williamson, Qian Zeng, Tricia Zwiefelhofer, Clinton R. Paden, Suxiang Tong, Duncan MacCannell |
| EPI_ISL_1364518 | Stanford Health Care | Stanford University School of Medicine, Clinical Virology Laboratory | Daniel Solis, Mamdouh Sibai, Fumiko Yamamoto, Malaya K. Sahoo, Michelle Verghese, ChunHong Huang, James Zehnder, and Benjamin A. Pinsky |
| EPI_ISL_593554, EPI_ISL_593555, EPI_ISL_593556, EPI_ISL_593557, EPI_ISL_593558 | Brigham and Women's Hospital | Jonathan Li Laboratory | Manish C. Choudhary, James Regan, Jonathan Z. Li |
| EPI_ISL_604182 | Quest Diagnostics | Quest Diagnostics | Rosenthal,S.H., Gerasimova,A., Kagan,R.M., Anderson, B., Grover, D., Livingston, K.E., Hua, M., Liu Y., Shalhout, D.F., Owen, R., Lacbawan, F. |
| EPI_ISL_852169, EPI_ISL_852237 | Lighthouse Lab in Milton Keynes | Wellcome Sanger Institute for the COVID-19 Genomics UK (COG-UK) Consortium | The Lighthouse Lab in Milton Keynes and Alex Alderton, Roberto Amato, Sonia Goncalves, Ewan Harrison, David K. Jackson, Ian Johnston, Dominic Kwiatkowski, Cordelia Langford, John Sillitoe on behalf of the Wellcome Sanger Institute COVID-19 Surveillance Team |
| EPI_ISL_876519 | DOHMH Jamaica | New York City Public Health Laboratory | Jade Wang, et al. |
| EPI_ISL_905171 | Dutch COVID-19 response team | National Institute for Public Health and the Environment (RIVM) | Adam Meijer, Harry Vennema, Dirk Eggink, Jeroen Cremer, Sharon van den Brink, Bas van der Veer, AnneMarie van den Brandt, Florian Zwagemaker, Dennis Schmitz, Chantal Reusken, on behalf of the national COVID-19 response team |
| EPI_ISL_955135 | Colorado Department of Public Health and Environment | Colorado Department of Puplic Health and Environment | Laura Bankers, Molly C. Hetherington-Rauth, Diana Ir, Shannon Ely, Shannon R. Matzinger, Sarah Elizabeth Totten, Emily A. Travanty |
| EPI_ISL_984758, EPI_ISL_984895, EPI_ISL_984920, EPI_ISL_994917, EPI_ISL_994936 | Pandemic Response Lab - NYC | Pandemic Response Lab, R&D | Henry Lee, Michael Hammerling, Melissa Hopkins, Cybill del Castillo, William Ward, Pradeep Bugga, Haiping Hao, Jon Laurent |
