## Supplementary material for "A new SARS-CoV-2 lineage that shares mutations with known Variants of Concern is rejected by automated sequence repository quality control": Figures S1 and S2, Tables S2 and S3, all supplemental figure and table legends

**Supplemental Tables and Figures:**

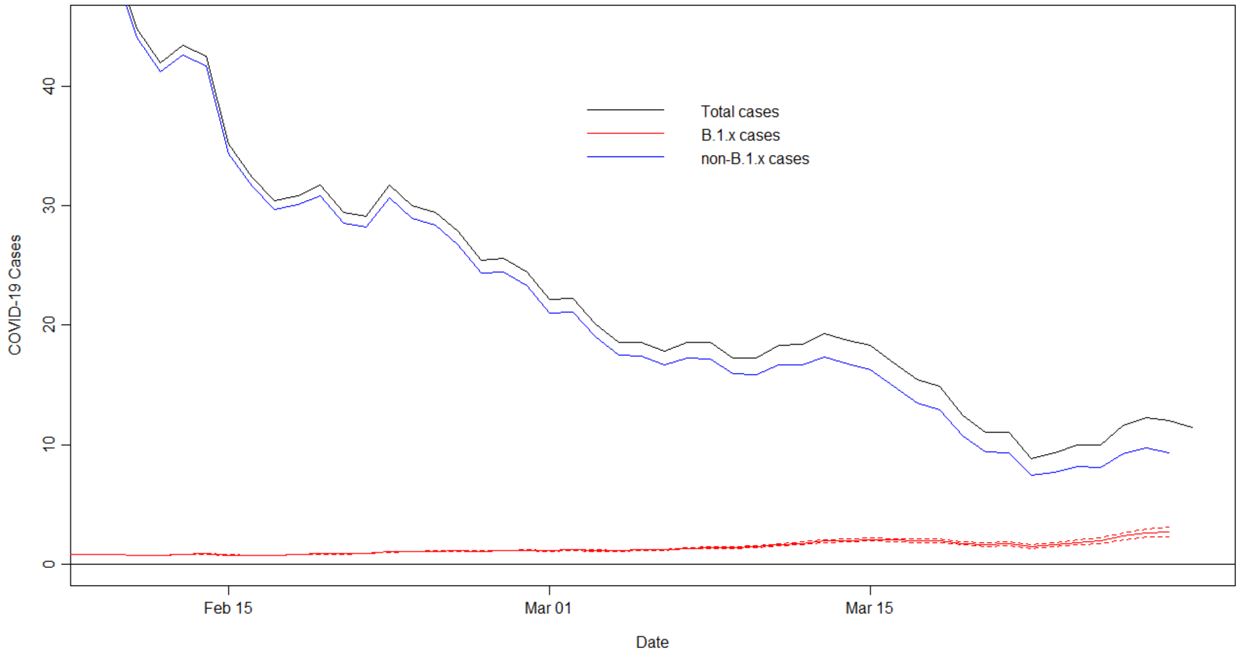


***Figure S1. Covid-19 cases in Santa Cruz County from Feb 10 to March 29, 2021, plotted as the 7-day running average.*** *The black line shows total cases. The red line shows B.1.x cases (with dashed lines showing the 95% CI) estimated by multiplying the total cases by the estimated fraction of samples that were B.1.x using the statistical model in Figure 2. The blue line shows the remaining non-B.1.x cases.*

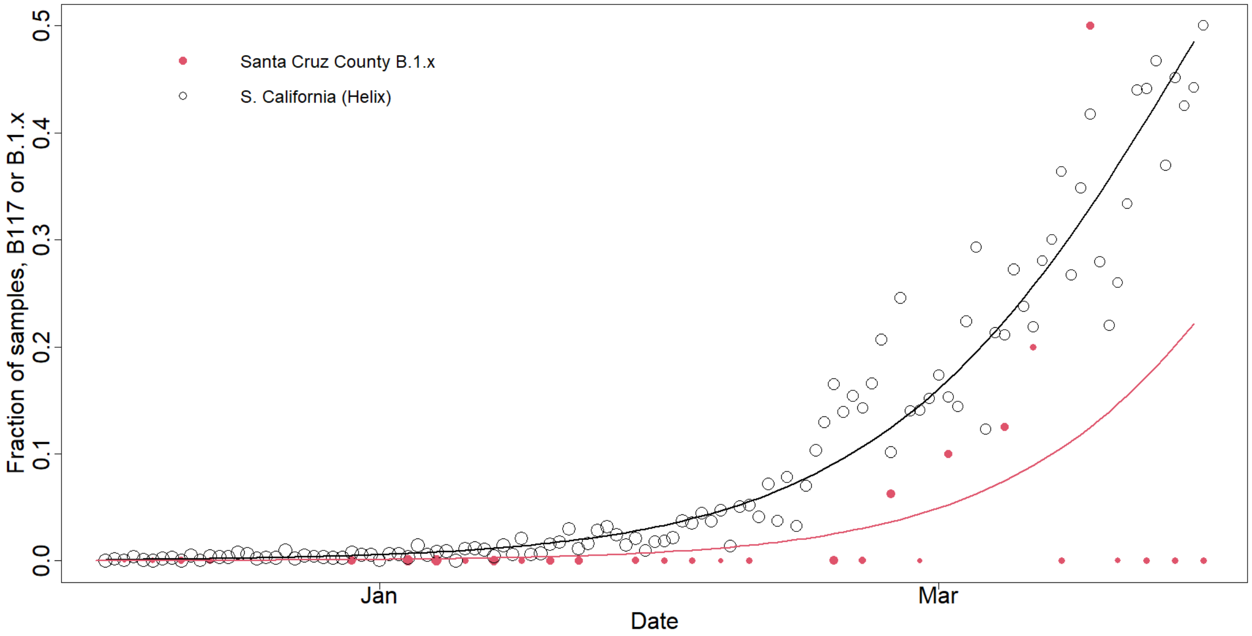


***Figure S2. Fraction of samples that are lineage B.1.1.7 (based on Helix data from southern California) or B.1.x in Santa Cruz County from December 3, 2020 through March 29, 2021.*** *Size of points shows the log-transformed relative sample size (Helix B.1.1.7 points: mean 493, range 40-1607; Santa Cruz County B.1.x points: mean 6.7, range 1-48). The fitted lines are based on generalized linear models with a binomial distribution (B.1.1.7: intercept -1104 ± 19.7; slope 0.059 ± 0.0011; P < 0.001; B.1.x: intercept -1176 ± 378; slope 0.063 ± 0.020; P = 0.002). A model fit to both time series with variant (B.1.x vs B.1.1.7; B.1.1.7 was the reference level) interacting with date indicated non-significant differences for initial frequency of B.1.x (intercept difference -71 ± 378; P = 0.85) and rate of increase for B.1.x (slope difference 0.0038 ± 0.020; P = 0.85).*

***Table S1:*** *UCSC sequence IDs, dates, clades/lineages. Many sequences are missing accession numbers in the table, especially for GenBank. Most sequences that don't yet have GenBank accessions will eventually, although a few sequences were rightfully rejected.*

| Pango lineage | Nucleotide mutation causing N:M234I |
|---|---|
| A.2.5 | G28975T |
| AB.1 | G28975C |
| B.1.1.214 | G28975T |
| B.1.1.397 | G28975T |
| B.1.1.445 | G28975C |
| B.1.1.486 | G28975T |
| B.1.36.21 | G28975T |
| B.1.160 | G28975C |
| B.1.328 | G28975T |
| B.1.460 | G28975T |
| B.1.523 | G28975T |
| B.1.526 | G28975A |
| B.1.561 | G28975T |
| B.1.585 | G28975T |
| B.1.588 | G28975T |
| P.2 | G28975T |

***Table S2:*** *Pango lineages with N:M234I in at least 80% of lineage-defining samples.*

| | Sequenced | Cases | Percent Sequenced |
|---|---|---|---|
| January | 143 | 4497 | 3.2 |
| February | 52 | 1109 | 4.7 |
| March | 52 | 428 | 12.1 |

***Table S3:*** *Summary of COVID-19 cases in Santa Cruz County in the past 3 months, compared to the number of cases that we sequenced.*

***Table S4****: All samples from figure 4 in the order that they appear in the tree from top to bottom.*

***Table S5****: GISAID Acknowledgments Table for all GISAID samples shown in Figure 4.*
